# Pharmacological rescue of the brain cortex phenotype of *Tbx1* mouse mutants: significance for 22q11.2 deletion syndrome

**DOI:** 10.1101/2021.02.04.429794

**Authors:** Ilaria Favicchia, Gemma Flore, Sara Cioffi, Gabriella Lania, Antonio Baldini, Elizabeth Illingworth

## Abstract

**Objectives:** *Tbx1* mutant mice are a widely used model of 22q11.2 deletion syndrome (22q11.2DS) because they manifest a broad spectrum of physical and behavioral abnormalities that is similar to that found in 22q11.2DS patients. In *Tbx1* mutants, brain abnormalities include changes in cortical cytoarchitecture, hypothesized to be caused by the precocious differentiation of cortical progenitors. The objectives of this research are to identify drugs that have efficacy against the brain phenotype, and through a phenotypic rescue approach, gain insights into the pathogenetic mechanisms underlying *Tbx1* haploinsufficiency.

**Experimental approach:** *Disease model:* *Tbx1* heterozygous and homozygous embryos. We tested the ability of two FDA-approved drugs, the LSD1 inhibitor Tranylcypromine and Vitamin B12, to rescue the *Tbx1* mutant cortical phenotype. Both drugs have proven efficacy against the cardiovascular phenotype, albeit at a much reduced level compared to the rescue achieved in the brain.

*Methods:* *in situ* hybridization and immunostaining of histological brain sections using a subset of molecular markers that label specific cortical regions or cell types. Appropriate quantification and statistical analysis of gene and protein expression were applied to identify cortical abnormalities and to determine the level of phenotypic rescue achieved.

**Results:** Cortical abnormalities observed in *Tbx1* mutant embryos were fully rescued by both drugs. Intriguingly, rescue was obtained with both drugs in *Tbx1* homozygous mutants, indicating that they function through mechanisms that do not depend upon *Tbx1* function. This was particularly surprising for Vitamin B12, which was identified through its ability to increase *Tbx1* gene expression.

**Conclusions:** To our knowledge, this is only the second example of drugs to be identified that ameliorate phenotypes caused by the mutation of a single gene from the 22q11.2 homologous region of the mouse genome. This one drug-one gene approach might be important because there is evidence that the brain phenotype in 22q11.2DS patients is multigenic in origin, unlike the physical phenotypes, which are overwhelmingly attributable to *Tbx1* haploinsufficiency. Therefore, effective treatments will likely involve the use of multiple drugs that are targeted to the function of specific genes within the deleted region.

## INTRODUCTION

Brain-related phenotypes, including intellectual disability and psychiatric disorders, are a major concern for the clinical management and quality of life of patients affected by 22q11.2 deletion syndrome (22q11.2DS). Several of the many genes heterozygously deleted in patients have been suggested to contribute to this group of phenotypes, based on data obtained in animal models. However, *TBX1*, the major candidate gene for the phenotypic findings linked to development of the embryonic pharynx, has been found to be mutated in patients with a 22q11.2DS-like phenotype that includes neurobehavioral anomalies. For example, *TBX1* mutation has been found in a patient with Asperger syndrome (Paylor et al., 2006) and in association with developmental delay (Ogata et al., 2014; Torres-Juan et al., 2007) and attention deficits (Haddad et al., 2019). Therefore, *TBX1* should be regarded as a major candidate for at least some of the brain-related clinical presentations in 22q11.2DS patients.

Mouse models have been instrumental not only for identifying the role of *Tbx1* in the disease, but also for determining its role in mouse behavior and brain development (Flore et al., 2017; Motahari et al., 2019; Paylor and Lindsay, 2006; Paylor et al., 2006; Prasad et al., 2008). *Tbx1* mutant mouse embryos have a complex brain cortex phenotype (Flore et al., 2017). Most interestingly, the heterozygous phenotype is almost as severe as the homozygous phenotype, indicating a high sensitivity of the brain cortex to *Tbx1* dosage. An additional peculiarity of Tbx1 function in the embryonic mouse brain is that it is mesoderm-dependent, thus mediated by an extracellular signal. This is also the case for several other phenotypes associated with the haploinsufficiency or loss of this gene (Braunstein et al., 2009; Calmont et al., 2009; Fagman et al., 2007; Lania et al., 2009; Xu et al., 2007; Zhang et al., 2005). In addition, recently, the mouse models have been used for testing experimental strategies for rescuing the cardiovascular phenotype, opening the way to targeted pre-clinical studies. The rescue of phenotypes caused by deficiency of *Tbx1*, which encodes a transcription factor, presents special challenges because of the potential number and variety of target genes. Nevertheless, research has identified a number of targeted strategies that have been successful in ameliorating the cardiovascular phenotype (Caprio and Baldini, 2014; Fulcoli et al., 2016; Lania et al., 2016; Racedo et al., 2017; Vitelli et al., 2010). However, the brain cortex phenotype has never been tested. Given the importance of the brain-related phenotype associated with 22q11.2DS, and because of the potential opportunity for clinical application, we tested two rescuing strategies in mice, selected on the basis of their potential clinical applicability. Results showed that the Lsd1/2 inhibitor Tranylcypromine (TCP) and Vitamin B12 (B12) prevent brain cortex abnormalities in *Tbx1* mutants.

## MATERIALS AND METHODS

### Mouse lines and in vivo drug treatments

Mouse studies were performed according to current regulations under the animal protocol 257/2015-PR approved by the Italian Ministry of Health. *Tbx1*^*lacz*/+^ (Lindsay et al., 2001) mice are maintained on a C57BL/6N background, *Tbx1*^+/Δ*e5*^ (Xu et al., 2004). Genotyping was performed as in the original reports. Administration of vitamin B12 (cyanocobalamin Sigma-Aldrich, V2876) was performed by intraperitoneal (i.p.) injection of pregnant female mice daily from E7.5 to E11.5 with 20mg Kg^−1^ body weight of B12 dissolved in PBS. Control mice were injected with the same volume of PBS. Administration of TCP was performed as described for vitamin B12, using 10mg Kg^−1^ body weight of TCP (Sigma-Aldrich, p8511-G) dissolved in sterile 0.9% w/v NaCl solution. Control mice were injected with the same volume of sterile 0.9% w/v NaCl solution. Developmental staging was establshed considering the morning of a visible vaginal plug as stage E0.5.

### Phenotype analysis

Pregnant females were sacrificed at E13.5. Phenotype evaluations were performed blind to the genotype.

#### a) *In situ* hybridization on mouse embryo brain sections

##### Cryosections

E13.5 brains were fixed in 4% PFA/1x PBS at 4°C overnight, washed in 1x PBS and incubated for 12 h in serial dilutions of sucrose/1x PBS (10%, 20%, 30% sucrose) at 4°C. Brains were then incubated for 2 h at 4°C in 50:50 v/v 30% sucrose/1x PBS/OCT prior to embedding in OCT. Twenty micron coronal sections were cut along the rostral–caudal brain axis on a cryostat and were postfixed in 4% PFA at RT for 10 min prior to immunostaining, except for immunofluorescence (IF). Alternatively, specimens were stored at −80°C. For *in situ* hybridization (ISH), antisense RNA probes were labeled using a DIG-RNA labeling kit (Roche). The probes used were: *Pax6*, *Tis21* and *NeuroD2*; DIG-labeled probes were hybridized to cryosections following published methods (Hirsch et al., 1998).

#### b) Immunofluorescence and immunohistochemistry on mouse embryo brain sections

Immunofluorescence (IF) was performed using the following primary antibodies: Mouse monoclonal anti-human KI67 antibody, 1:200 (BD Pharmingen, #556003) and rabbit polyclonal anti-PH3 antibody, 1:200 (Millipore, #06-570). Experiments were performed on serial sections 200 μm apart, using a minimum of 5 embryos per genotype. Sections were briefly microwaved to boiling point in 10 mM sodium citrate (pH 6.0) 3 times, for antigen enhancement, cooled, rinsed in 1x PBS, permeablized with 0.1% Triton x-100 blocked in 10% GS/PBS/0.1% Triton x-100 for at least 1h at RT. Sections were then incubated with primary antibodies overnight at 4°C in the same blocking solution reducing the GS to 5%, rinsed, and incubated in secondary antibodies for at least 1h 30 min at RT. The following secondary antibodies were used: For PH3, Alexa Fluor 594-conjugated anti-rabbit, 1:400 (Life Technologies, Invitrogen). For anti-KI67 immunohistochemistry (IHC), sections were also treated with 0.5% H2O2 in ethanol to block endogenous peroxidase activity. Anti-KI67, was revealed by incubation with a biotinylated mouse secondary antibody (1:200) and VECTASTAIN Elite-ABC kit reaction (Vector Laboratories) with the TSA-Plus Fluorescein System NEL741001KT (Perkin-Elmer Inc.), following the manufacturer’s instructions. Fluorescence was observed with an epifluorescence microscope (Leica DMI6000B, acquisition software LAS AF 2.6). Images were digitally documented with a camera and computer processed using Adobe Photoshop® version 6 for Windows.

Immunohistochemistry (IHC) experiments were performed on serial sections 200 μm apart, using a minimum of 5 embryos per genotype. Sections were briefly post-fixed with 4% PFA in PBS, then rinsed with PBS, permeablized with 0.5% Tween 20, blocked in 20% GS/PBS/0.5% Tween 20 for at least 1h at RT. Sections were then incubated with the primary antibody (rabbit polyclonal anti-TBR1 antibody, 1:200, Millipore, #AB10554) overnight at 4°C in the same blocking solution reducing the GS to 5%, rinsed, and treated with 0.5% H_2_O_2_ in ethanol to block endogenous peroxidase activity. TBR1-expressing cells were revealed by incubation with biotinylated rabbit secondary antibody (1:200), VECTASTAIN Elite-ABC kit reaction and SK-4100 DAB Substrate Kit (Vector Laboratories), following the manufacturer’s instructions. Staining was observed in bright field with a Leica DM6000B microscope (acquisition software LAS V4.1). Images were digitally documented with a camera and computer processed using Adobe Photoshop® version 6 for Windows.

#### c) Data quantification

Cell counts on E13.5 brain sections (IF and IHC): For PH3 quantification, cell counts were performed on a single hemisphere per embryo (n = 5–6/genotype), comprising nine anatomically matched sections 200 μm apart along the rostral to caudal brain axis. Landmarks: the first 5 sections were 200 μm apart, starting from the first section in which the LGE was clearly visible, and before the appearance of the hippocampal primordium. In the mouse strain used here, this position is approx. 600 μm caudal to the prospective olfactory bulb (OB). Counting box positioning: in the cortex (Cx), multiple counting boxes of 100 μm^2^ spanning the thickness of the Cx, starting from the ventricular surface were positioned in the medio-lateral (ML) cortex, along a line tangential to the LGE or caudal (C) GE (cartoon in Fig. 1). For KI67 quantification, the ratio between the thickness of the Ki67-positive zone and the whole cortical thickness, in the ML cortex, was evaluated on a single hemisphere per embryo (n = 5–6/genotype). Seven sections per hemisphere were evaluated in rostral - caudal sections following the aforementioned landmarks. The 6^th^ and 7^th^ sections were 400 μm apart, so as to identify better the CGE domain. For TBR1 quantification, sections were processed following the landmarks indicated for *ISH* analysis, and TBR1-positive cells were counted using multiple counting boxes of 100 μm^2^ spanning the thickness of the Cx, positioned starting from the ventricular surface in the medio-lateral (ML) cortex, along a line tangential to the GEs (lateral, medial or caudal at the corresponding levels).

**FIGURE 1.**
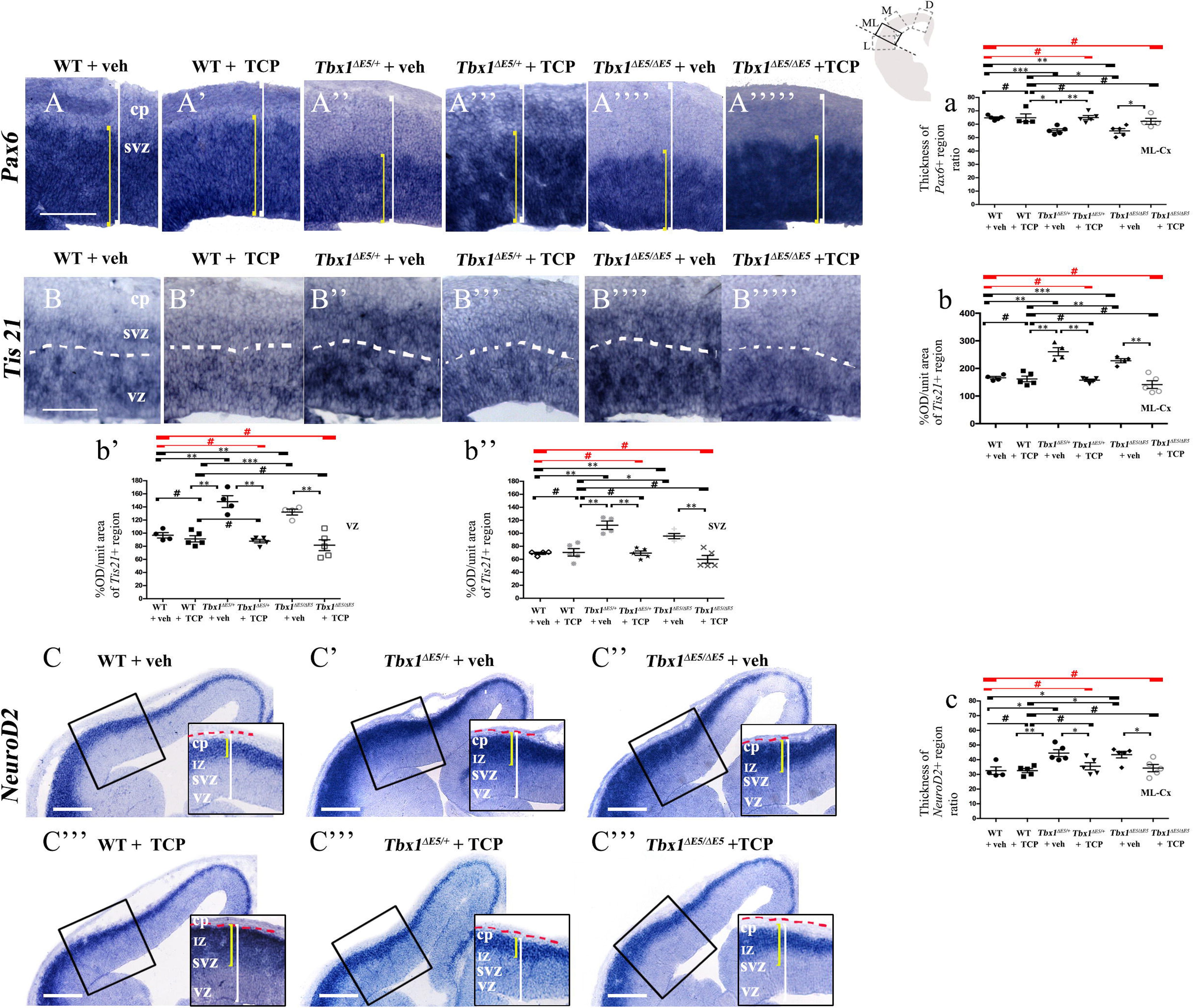
TCP treatment restores normal expression of cortical markers in *Tbx1* mutant embryos. Panels show coronal sections of embryonic brains at E13.5. The cartoon in A indicates the position of the ML-cortex in coronal brain sections, where all measurements were made. (A, a) *In situ* hybridization of *Pax6* (A–A’′′”) revealed thinning of the expression domain (yellow brackets) in vehicle-treated *Tbx1*^*ΔE5*/+^ (heterozygous) embyos (A”) and *Tbx1*^*ΔE5/ΔE5*^ (homozygous) embyos (A””) compared to vehicle-treated WT embryos (A, a), which was rescued by TCP treatment (A”’, A””’, a). (B, b) *In situ* hybridization of *Tis21* (B–B′′”’) showed increased expression (OD) in the ML-cortex of vehicle-treated *Tbx1*^*ΔE5*/+^ embryos (B”, b) and *Tbx1*^*ΔE5/ΔE5*^ embyos (B””, b), compared to vehicle-treated WT embryos (B, b), including the VZ (b’) and SVZ (b”), which was rescued by TCP treatment (B”’, B””’, b, b’, b”). (C, c) *In situ* hybridization of *NeuroD2* (C–C′′”’) showed increased expression in the ML-cortex of vehicle-treated *Tbx1*^*ΔE5*/+^ embryos (C′, c) and *Tbx1*^*ΔE5/ΔE5*^ embyos (C”, c) compared to vehicle-treated WT embryos (C, c), and extension of the expression domain (yellow brackets in C’, C’′) relative to the total ventricular pial thickness (white brackets), which was rescued by TCP treatment (C””, C””’, c).

##### *In situ* evaluation

For *Tis21*, optical density (OD), minus the slice background OD, was measured using the specific ImageJ tool (unit area 50 μm^2^), while for Pax6 and ND2, we measured the ratio between the thickness of the probe signal and the total thickness of the ML cortex, using the ImageJ graphic pen on a single hemisphere in rostral, medial and caudal positions that were 600 μm apart. Landmarks in coronal sections, rostral - caudal: Rostral: regarding the serial slices collected on each slide, the rostral slice selected was the second in which the LGE was clearly visible, before the appearance of the hippocampal primordium (approx. 800 μm after the appearance of the prospective OB). Medial: approx. 600 μm caudal to the rostral slice. The slice selected was the first in which the MGE was completely visible, together with the hippocampal primordium detached from the subpallium, the choroid plexus within the left and lateral ventricle and arising from the medial wall. Caudal: approx. 600 μm caudal to the medial slice. The slice was selected where the CGE was the only GE present. Landmarks for this level are the presence of the thalamus, superior and inferior part of 3^rd^ ventricle, interthalamic adhesion.

### Statistical Analysis

The data were statistically analyzed and graphically represented using the Microsoft Office Excel and Prism software. Results are expressed as the mean ± standard error of the mean (SEM). The unpaired Students *t*-test with Welch’s correction and Ordinary One Way ANOVA, were used for statistical analysis. For all experiments, values of P ≤ 0.05 were considered to be statistically significant.

## RESULTS

### LSD1/2 inhibition

Tranylcypromine (TCP) is an LSD1/2 (KDM1A/B) demethylase inhibitor. It acts by covalently binding to the coenzyme FAD, forming a covalent adduct with the flavin ring, thereby blocking irreversibly the enzymatic activity of LSD1 (Binda et al., 2002; Forneris et al., 2005; Shi et al., 2004). TCP has been shown to rescue partially the loss of H3K4me1 in *Tbx1*^*lacZ*/+^ embryos and partially rescue the cardiovascular phenotype in *Tbx1*^*lacZ*/+^ and *Tbx1*^*neo2*/−^ embryos (Fulcoli et al., 2016). To test whether TCP rescues or modifies the brain cortex phenotype in *Tbx1* mutants, we intercrossed *Tbx1*^Δ*e5*/+^ animals and pregnant females were injected intraperitoneally (i.p.) with TCP (10mg Kg^−1^ body weight) or with vehicle (saline solution) at embryonic days (E)10.5, E11.5, and E12.5. We harvested embryos at E13.5 and prepared coronal brain cryosections for marker analyses. Our previous study has shown that *Tbx1* mutants (heterozygous and homozygous loss of function) have brain cortex abnormalities that are consistent with premature differentiation of cortical neurons (Flore et al., 2017). In this study, we evaluated phenotypic rescue based on the expression of a subset of the molecular markers used by Flore et al., 2017, including, *Pax6*, *Tis21*, and *NeuroD2*. *Pax6* labels apical neural progenitors (AP) in the ventricular zone (VZ) of the cortex; its expression domain was thinned in *Tbx1* mutant embryos. *Tis21* identifies neurogenic progenitors, i.e., APs switching from symmetric self-renewing divisions to neurogenic divisions, and basal neural progenitors (BP) undergoing neurogenic divisions. Expression of *Tis21* was increased in *Tbx1* mutant embryos. *NeuroD2* labels fully differentiated neurons; its expression domain is expanded in *Tbx1* mutant embryos (Flore et al., 2017). Analysis of these markers in *Tbx1* mutant embryos confirmed these previously reported expression anomalies and showed that TCP treatment fully rescued their expression in *Tbx1^+/^*^Δ*e5*^ and *Tbx1*^Δ*e5/*Δ*e5*^ embryos, i.e., their expression returned to WT levels (Fig. 1A, 1a, *Pax6*; 1B, 1b, *Tis21*; 1C, 1c, *NeuroD2*, and Table 1). Moreover, WT embryos treated with vehicle or with TCP showed no differences in the cortical expression of *Pax6, Tis21* and *NeuroD2*, indicating that TCP alone does not influence cortical development (Fig. 1a, *Pax6*; 1b, *Tis21*; 1c, *NeuroD2*, and Table 1). Finally, *Tbx1*^Δ*e5*/+^ and *Tbx1*^Δ*e5/*Δ*e5*^ embryos treated with vehicle showed similar alterations in expression of *Pax6, Tis21* and *NeuroD2* as those previously reported by Flore et al., 2017 (Fig. 1a, *Pax6*; 1b, *Tis21*; 1c, *NeuroD2*, and Table 1).

**Table 1.**

| Unpaired t-test with Welch's Correction | WT-veh vs WT -TCP | WT-veh vs *Tbx1*<sup>-E5/+</sup>-veh | WT-TCP vs *Tbx1*<sup>-E5/+</sup>-TCP | WT-veh vs *Tbx1*<sup>-E5/+</sup>-TCP | WT-veh vs *Tbx1*<sup>-E5/-E5</sup>-veh | WT-TCP vs *Tbx1*<sup>-E5/-E5</sup>-TCP | WT-veh vs *Tbx1*<sup>-E5/-E5</sup>-TCP | Ordinary One Way Anova |
| --- | --- | --- | --- | --- | --- | --- | --- | --- |
| *Pax6* | 0.9886 | 0.0005*** | 0.9653 | 0.9130 | 0.0026** | 0.4970 | 0.3394 | 0.0002*** |
| *Tis21* | 0.4105(vz),<br>0.8255 (svz)<br>0.7327 (tot) | 0.0057** (vz),<br>0.0050** (svz)<br>0.0048** (tot) | 0.5466 (vz),<br>0.8697 (svz)<br>0.6949 (tot) | 0.1218 (vz)<br>0.9477 (svz)<br>0.2333 (tot) | 0.0011** (vz),<br>0.0037** (svz)<br>0.0009*** (tot) | 0.3476 (vz),<br>0.2341 (svz)<br>0.2471 (tot) | 0.1546 (vz),<br>0.1960 (svz)<br>0.1604 (tot) | <0.0001****<br>(vz+svz)<br>< 0.0001****<br>(tot) |
| *NeuroD2* | 0.9948 | 0.0116* | 0.3317 | 0.4322 | 0.0163* | 0.5597 | 0.6419 | 0.0035** |

### Vitamin B12 treatment

Vitamin B12 (B12) was identified in a high throughput screen (HTS) to identify small molecules that enhanced *Tbx1* gene expression in *Tbx1* heterozygous mouse embryonic fibroblasts (Lania et al., 2016). The authors of this study reported the partial rescue of cardiovascular phenotypes in mid- and pre-term *Tbx1* mutant embryos when treated with high doses of B12.

B12 is an essential vitamin that plays an important role in cellular metabolism through its effects on DNA synthesis, methylation and mitochondrial metabolism. It is activated in lysosomes through the formation of Methyl-B12 and Adenosyl-B12, which act as cofactors for methionine synthase and methylmalonyl-CoA mutase respectively. Methionine synthase catalyzes the conversion of Homocysteine into Methionine, which is then converted to S-adenosyl-methionine (SAM), a methyl donor for methyltrasferases, enzymes that modify the methylation status of histones and DNA. Methylmalonyl-CoA mutase is a mitochondrial enzyme that converts methylmalonyl-CoA into Succinyl-CoA, an intermediate of the citric acid cycle that is important for fatty acid and amino acid metabolism. Cases of clinically severe B12 deficiency, such as pernicious anemia, have revealed the critical role of this vitamin for hemopoiesis and for brain development and function (Green et al., 2017).

To test whether B12 treatment rescues or modifies the brain cortex phenotype in *Tbx1* mutants, we crossed male *Tbx1*^*lacZ*/+^ mice with female WT mice and pregnant females were injected i.p. with B12 (20mg/Kg^−1^ body weight) or with the same volume of vehicle (PBS) daily from E7.5 to E11.5. We harvested embryos at E13.5 and prepared coronal brain cryosections which were subjected to ISH with probes against *Pax6, Tis21* and *NeuroD2*, and immunostaining with anti-TBR1 antibody, which labels terminally differentiated cortical neurons. The results obtained with these markers, which are shown in Figs. 2 and 3 and summarized in Table 2, were similar overall to those obtained with TCP treatment. Specifically, in *Tbx1*^*lacZ*/+^ embryos, B12 treatment fully rescued the cortical expression of *Pax6, Tis21* and *NeuroD2*, compared to those treated with vehicle (Fig. 2A’, 2A”’, 2a’, *Pax6*; 2B’, 2B”’, 2b, 2b’, *Tis21*; 2C’, 2C”’, 2c, *NeuroD2* and Table 2). Moreover, WT embryos treated with the vehicle or with B12 showed no differences in cortical expression of these markers, indicating that B12 alone does not affect cortical development (Fig. 2A, 2A”, 2a’, 2b, *Pax6*; 2B, 2B”, 2b, 2b’, *Tis21*; 2C, 2C”, 2c, *NeuroD2* and Table 2). Finally, *Tbx1*^*lacZ*/+^ embryos treated with vehicle showed similar alterations in expression of *Pax6, Tis21*, *NeuroD2* as those previously reported by Flore et al., 2017 (Fig. 2A’, 2a, *Pax6*; 2B’, 2b, 2b’, *Tis21*; 2C’, 2c, *NeuroD2* and Table 2). Expression of TBR1 was also normalized in B12 treated *Tbx1*^*lacZ*/+^ embryos compared to those treated with vehicle (Fig. 3A’, 3A”’, 3a, 3a’).

**FIGURE 2.**
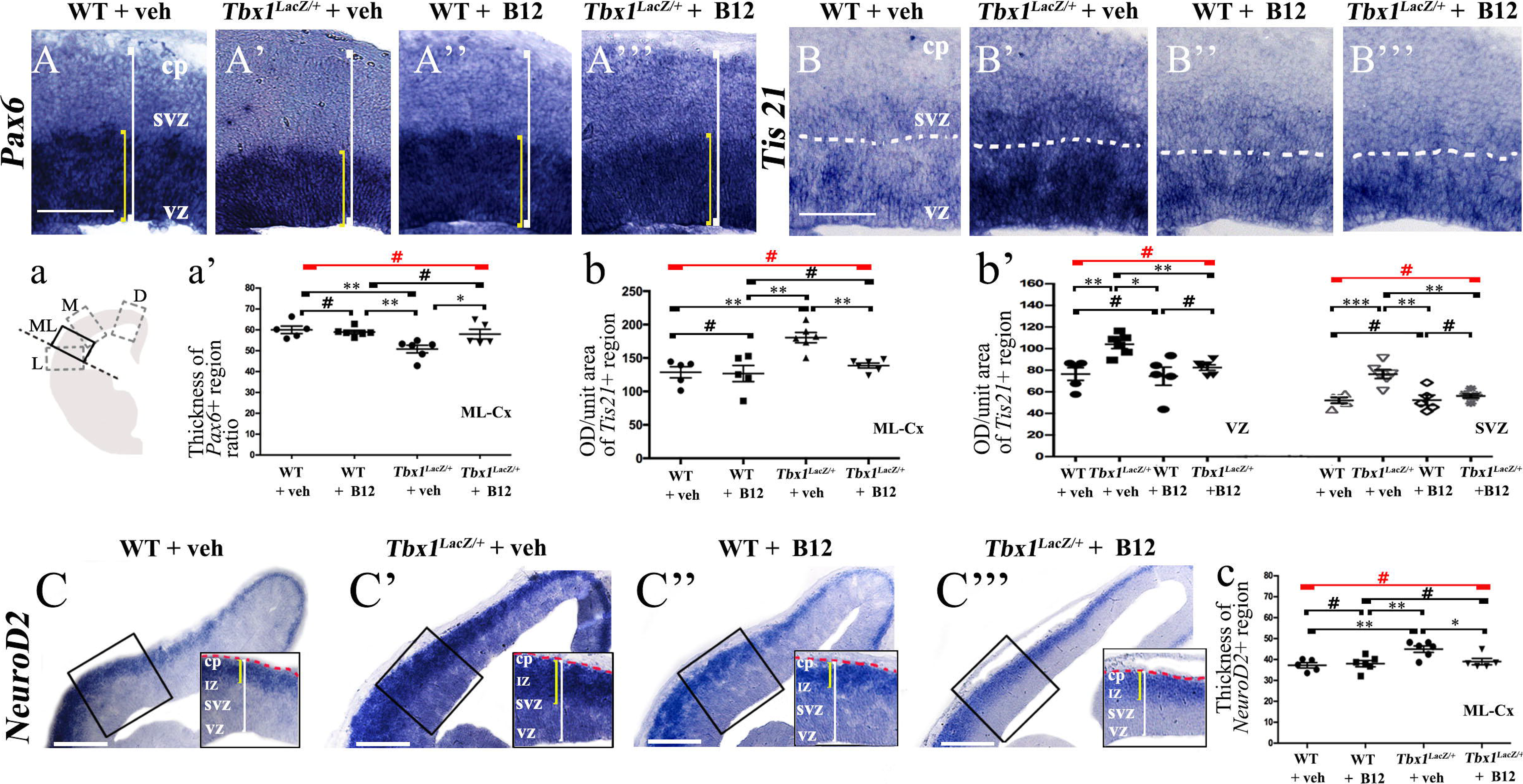
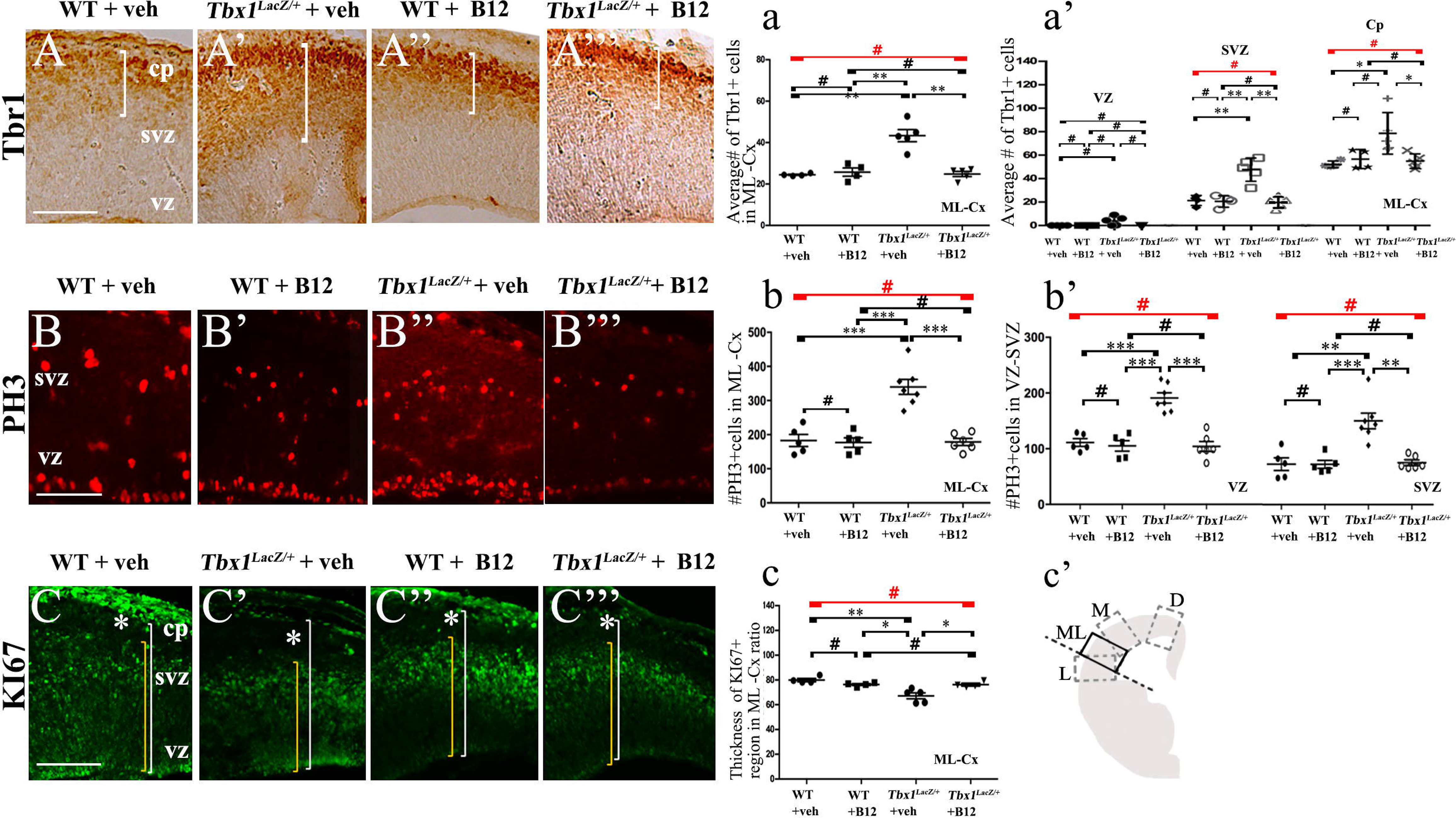
Vitamin B12 treatment restores normal expression of cortical markers in *Tbx1* heterozygous embryos. Panels show coronal sections of embryonic brains at E13.5. The cartoon (a) indicates the position of the ML-cortex in coronal sections, where all measurements were made. A, a) *In situ* hybridization of *Pax6* (A–A’′′) revealed thinning of the expression domain (yellow brackets) in vehicle-treated *Tbx1*^*lacZ*/+^ (heterozygous) embryos (A’) compared to vehicle-treated WT embryos (A, a), which was rescued by B12 treatment (A”’, a). (B, b, b’) *In situ* hybridization of *Tis21* (B–B′′’) showed increased expression (OD) in the ML-cortex of vehicle-treated *Tbx1^lacZ/+^* embryos compared to vehicle-treated WT embryos (B), including the VZ and SVZ (B’, b, b’), which was rescued by B12 treatment (B”’, b, b’). (C, c) *In situ* hybridization of *NeuroD2* (C–C′′’) showed increased expression in the ML-cortex of vehicle-treat *Tbx1*^*lacZ*/+^ embryos (C′, c) compared to vehicle-treated WT embryos (C) and extension of the expression domain (yellow brackets), relative to the total ventricular pial thickness (white brackets), which was rescued by B12 treatment (C”’, c). Dashed red line defines the external edge of the cortical plate (CP). Scale bars, 100 μm in all panels except panel C, which is 200 μm. *P ≤ 0.05, **P ≤ 0.01, ***P ≤ 0.001, error bars indicate ±SEM, n = 5–6 per genotype. Abbreviations: VZ, ventricular zone; SVZ, subventricular zone; CP, cortical plate; ML-Cx, medio-lateral cortex; D, dorsal; M, medial; ML, medio-lateral; L, lateral; OD, optical density.

**FIGURE 3.**
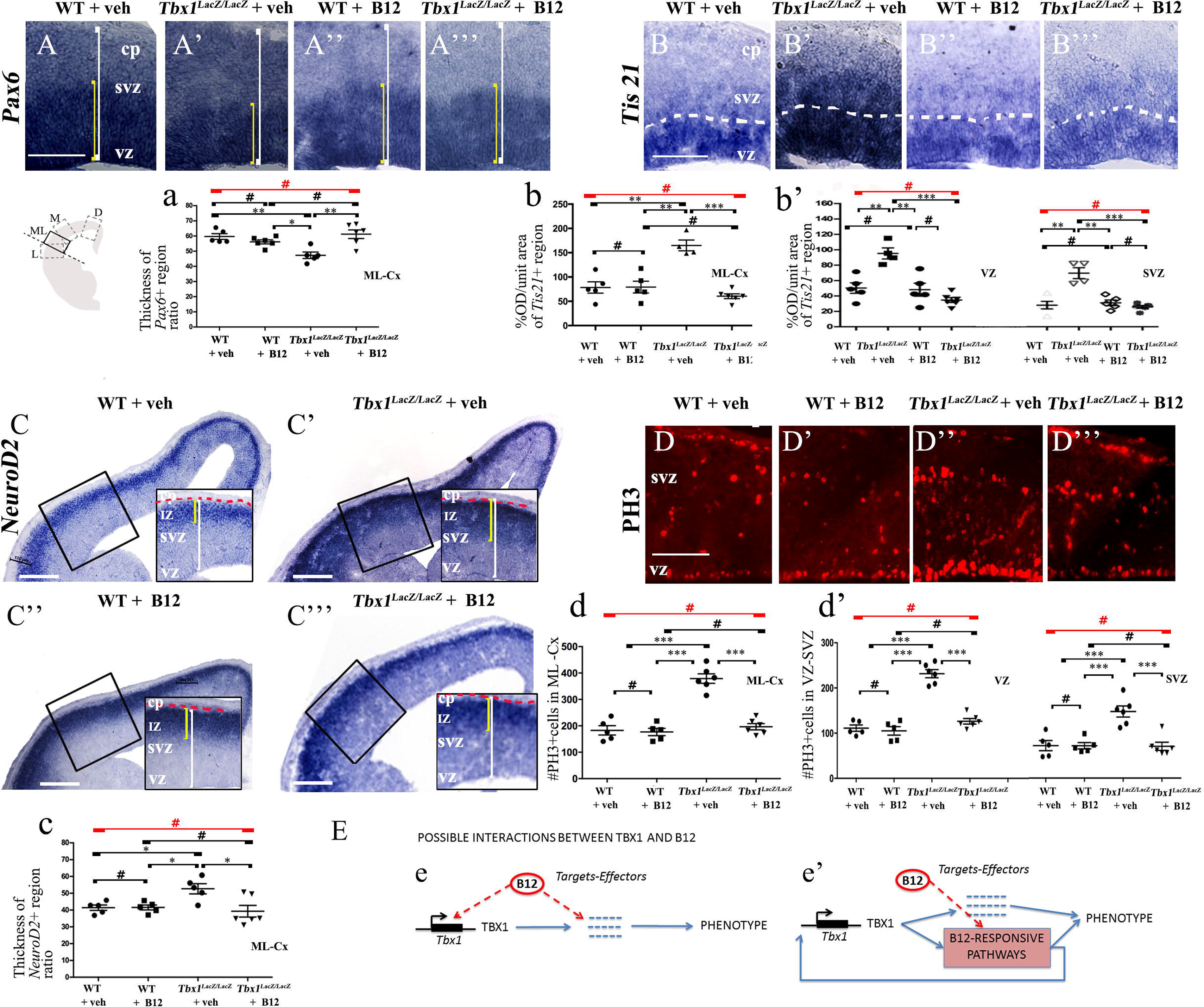
Vitamin B12 treatment restores normal expression of cortical markers in *Tbx1* heterozygous embryos. Panels show coronal sections of embryonic brains at E13.5. (A, a) Immunostaining for TBR1 (A–A′′’) revealed the precocious presence of terminally differentiated neurons in the SVZ of *Tbx1*^*lacZ*/+^ embryos (A’). White brackets define the regions containing TBR1-positive cells in A, A’. Quantitative analysis confirmed the altered distribution and frequency of mature cortical neurons in the ML-cortex, of *Tbx1*^*lacZ*/+^ embryos, including the SVZ and CP (a), which was rescued by B12 treatment (A”’, a). (B, b) Immunostaining for phospho-histone 3 (PH3) revealed increased mitotic activity in the proliferative zones (VZ, SVZ) of the ML-cortex of vehicle-treated *Tbx1*^*lacZ*/+^ embryos (B’, b, b’) compared to vehicle-treated WT embryos (B, b), which was rescued by B12 treatment (B”’, b, b’). (C, c) Immunostaining for KI67. White asterisks identify KI67-negative regions. Quantitative analysis (c) revealed the reduced thickness of the KI67-expressing region (yellow brackets) relative to the total ventricular-to-pial thickness (white brackets) in vehicle-treated *Tbx1*^*lacZ*/+^ embryos (C’, c) compared to vehicle-treated WT embryos (C), which was rescued by B12 treatment (C”’, c). The cartoon (c’) indicates the position of the ML-cortex in coronal sections, where all measurements were made. Scale bars, 100 μm. *P ≤ 0.05, **P ≤ 0.01, ***P ≤ 0.001, error bars indicate ±SEM, n = 5–6 per genotype. Abbreviations: VZ, ventricular zone; SVZ, subventricular zone; CP, cortical plate; ML-Cx, medio-lateral cortex; D, dorsal; M, medial; ML, medio-lateral; L, lateral; OD, optical density.

**Table 2.**

| Unpaired t-test with Welch's Correction | WT-veh vs WT B12 | WT-veh vs *Tbx1*<sup>LacZ/+</sup>-veh | WT-B12 vs *Tbx1*<sup>LacZ/+</sup>-B12 | WT-veh vs *Tbx1*<sup>LacZ/+</sup>-B12 | Ordinary One Way Anova |
| --- | --- | --- | --- | --- | --- |
| *Pax6* | 0.6073 | 0.0059** | 0.7118 | 0.5061 | 0.0047** |
| *Tis21* | 0.8485 (vz),<br>0.9754 (svz)<br>0.9030 (tot) | 0.0056** (vz),<br>0.0009*** (svz)<br>0.0015** (tot) | 0.4040 (vz),<br>0.4688 (svz)<br>0.3908 (tot) | 0.3889 (vz),<br>0.2299 (svz)<br>0.3134 (tot) | 0.0029** (vz)<br>0.0002 (svz)<br>0.0004*** (tot) |
| *NeuroD2* | 0.7050 | 0.0037** | 0.6428 | 0.3978 | 0.0047** |
| *PH3* | 0.6280(vz),<br>0.9884 (svz)<br>0.7930 (tot) | <0.0001**** (vz),<br>0.0015** (svz)<br>0.0002*** (tot) | 0.9342 (vz),<br>0.7426 (svz)<br>0.9194 (tot) | 0.511 (vz),<br>0.8316 (svz)<br>0.08401 (tot) | <0.0001**** (vz),<br><0.0001**** (svz)<br><0.0001**** (tot) |
| *Ki67* | 0.0764 | 0.0033** | 0.9379 | 0.0867 | 0.0007*** |
| *Tbr1* | 0.8256 (svz)<br>0.3504(cp)<br>0.5680(tot) | 0.0021** (svz)<br>0.0278* (cp)<br>0.0029** (tot) | 0.7952 (svz)<br>0.7296 (cp)<br>0.7193 (tot) | 0.5992 (svz)<br>0.4016 (cp)<br>0.7648 (tot) | <0.0001**** (svz)<br>0.0073** (cp)<br><0.0001**** (tot) |

In order to obtain a broader picture of the efficacy of B12 treatment, we looked for rescue of the cell proliferation/cell cycle anomalies, which are part of the cortical phenotype observed in *Tbx1* mutants (Flore et al., 2017). For this, we immunostained brain cryosections for phospho-histone 3 (PH3), which labels mitotic cells (M phase) and KI67, which labels cycling cells, i.e., not in G0. Flore et al. reported an increased number of PH3+ cells in the proliferating layers of the cortex in *Tbx1* mutants at E13.5, and an expanded KI67-negative layer where terminally differentiated neurons reside. We found that B12 treatment fully rescued both of these cellular phenotypes in the cortex of *Tbx1*^*lacz*/+^ embryos (Fig. 3B’, 3B”’, 3b, 3b’, PH3; 3C’, 3C”’, 3c, KI67 and Table 2). Together, these results indicate that B12 treatment has high efficacy in the developing murine cortex.

In order to understand whether B12 treatment restores normal gene expression and normal cortical cytoarchitecture through a mechanism that does not depend upon the transcriptional increase of *Tbx1*, we intercrossed *Tbx1*^*lacZ*/+^ mice and treated pregnant females with B12 using the same experimental conditions as those described for *Tbx1* heterozygous embryos. Surprisingly, we found that in B12-treated *Tbx1^lacZ/lacZ^* embryos, the cortical expression of *Pax6, Tis21*, *NeuroD2* and PH3 was fully rescued, compared to *Tbx1^lacZ/lacZ^* embryos treated with the vehicle (Fig. 4A’, 4A”’, 4a, *Pax6*; 4B’, 4B”’, 4b, 4b’, *Tis21*; 4C’, 4C”’, 4c, *NeuroD2*; 4D’, 4D”’, 4d, 4d’, PH3 and Table 3). These results demonstrate that the phenotypic rescue obtained with B12 does not depend solely upon the transcriptional increase of *Tbx1*. Figure 4E shows a model of the possible interactions between TBX1 and B12 that could account for the phenotypic rescue.

**Table 3.**

| Unpaired t-test with Welch's Correction | WT-veh vs WT B12 | WT-veh vs *Tbx1*<sup>LacZ/LacZ</sup>-veh | WT-B12 vs *Tbx1*<sup>LacZ/LacZ</sup>-B12 | WT-veh vs *Tbx1*<sup>LacZ/LacZ</sup>-B12 | Ordinary One Way Anova |
| --- | --- | --- | --- | --- | --- |
| *Pax6* | 0.1739 | 0.0027** | 0.1513 | 0.6634 | 0.0014** |
| *Tis21* | 0.8510 (vz),<br>0.6412 (svz)<br>0.9600 (tot) | 0.0030** (vz),<br>0.0036** (svz)<br>0.0011** (tot) | 0.1949 (vz),<br>0.2546 (svz),<br>0.2054 (tot) | 0.0871 (vz),<br>0.6768 (svz),<br>0.2086 (tot) | <0.0001**** (vz)<br><0.0001**** (svz)<br><0.0001**** (tot) |
| *NeuroD2* | 0.9544 | 0.0169* | 0.5735 | 0.5996 | 0.0111* |
| *PH3* | 0.6280 (vz),<br>0.9884 (svz)<br>0.7930 (tot) | <0.0001**** (vz),<br>0.0014** (svz)<br><0.0001**** (tot) | 0.0995 (vz),<br>0.9148 (svz)<br>0.3048 (tot) | 0.1355 (vz),<br>0.9214 (svz)<br>0.5365 (tot) | <0.0001**** (vz),<br><0.0001**** (svz)<br><0.0001**** (tot) |

**FIGURE 4.** Vitamin B12 treatment restores normal expression of cortical markers in *Tbx1* homozygous embryos. All panels show coronal sections of embryonic brains at E13.5. The cartoon in A indicates the position of the ML-cortex in coronal brain sections, where all measurements were made. (A, a) *In situ* hybridization of *Pax6* (A–A’′′) revealed thinning of the expression domain (yellow brackets) in vehicle-treated *Tbx1^lacZ/lacZ^* embryos (A’) compared to WT (A, a), which was rescued by B12 treatment (A”’, a). (B, b, b’) *In situ* hybridization of *Tis21* (B–B′′’) showed increased expression in the ML-cortex of *Tbx1^lacZ/lacZ^* embryos, including the VZ and SVZ (B’, b, b’), which was rescued by B12 treatment (B”’, b, b’). Dashed white line indicates border between the VZ and SVZ. (C, c) *In situ* hybridization of *NeuroD2* (C–C′′’) showed increased expression in the ML-cortex of *Tbx1^lacZ/lacZ^* embryos (C′, c) and extension of the expression domain (yellow brackets) relative to the total ventricular pial thickness (white brackets), which was rescued by B12 treatment (C”’, c). Dashed red line defines the external edge of the CP. (D, d, d’) Immunostaining for phosphohistone 3 (PH3) revealed increased mitotic activity in the proliferative zones (VZ, SVZ) of the ML-cortex of *Tbx1^lacZ/lacZ^* embryos (D’, d, d’), which was rescued by B12 treatment (D”’, d, d’). (E) Alternative models of interactions between TBX1 and B12 that are consistent with the observed phenotypic rescue in *Tbx1* homozygous mutants. In e, B12 acts by altering the expression of genes, including *Tbx1,* and one or more critical TBX1 targets. The latter activity is retained when *Tbx1* expression is abolished. In e’, B12 acts downstream of TBX1, on one or more critical pathways that respond to TBX1 and has a positive feedback on *Tbx1* expression. In the absence of TBX1, B12 continues to affect (activate or repress) the critical pathways.

## DISCUSSION

Our study shows for the first time that anomalies of cortical cytoarchitecture caused by *Tbx1* mutation can be prevented by appropriate pharmacological intervention during gestation. We tested two compounds that have been shown to reduce the frequency and severity of cardiovascular defects in *Tbx1* mutant mice, namely TCP and B12. We found that treatment with either compound fully restored the normal expression of a set of cortical markers that is anomalously expressed in *Tbx1* mutant embryos. We focused our attention on the rescue of the phenotypes affecting cortical progenitors (APs and BPs) and their progeny that were described by Flore et al., 2017. The study by Flore et al. showed that the cortical phenotype was fully penetrant and had a similar severity in *Tbx1*^*lacZ*/+^ and *Tbx1^lacZ/lacZ^* embryos at E13.5. In the current study, we found that this was also the case for a second *Tbx1* allele, *Tbx1*^Δ*e5*^ which is a null allele resulting from the recombination of the *Tbx1*^*flox*^ allele (Xu et al., 2004). Furthermore, for both alleles, and for both drugs, the phenotypic rescue was complete, for the markers tested. This is in contrast to the cardiovascular phenotype, for which only partial rescue was obtained (Fulcoli et al., 2016; Lania et al., 2016). Thus, the brain seems to be more sensitive than the heart to the beneficial effects of these two drugs. In the study by Fulcoli et al., only about one third of the genes, the expression of which was disrupted in cells with *Tbx1* knockdown, were rescued by TCP treatment. This might account, at least in part, for the incomplete phenotypic rescue obtained. In the cortex, the greater phenotypic rescue obtained with TCP (and with B12) might be due to a greater recovery of target gene expression, or TBX1 might have different critical targets in brain that are more sensitive to these drugs. Currently, the identity of the Tbx1-dependent cell population that is critical for cortical development in unknown, other than it is mesodermal in origin (Flore et al., 2017).

The rescue of the cortical phenotype in *Tbx1* homozygous mutants treated with TCP was not altogether surprising, since TCP is predicted to function downstream of TBX1 by reducing the H3K4me1 depletion caused by *Tbx1* loss of function (Fulcoli et al., 2016) i.e., through a non-transcriptional mechanism. However, the full phenotypic rescue obtained with B12 treatment was quite unexpected, given that B12 was identified for its ability to enhance *Tbx1* expression. Thus, our study indicates that B12, in addition to acting by transcriptionally increasing *Tbx1* expression, also acts by other as yet unknown mechanisms that favor phenotypic rescue. What might those mechanisms be? One possibility is that in the absence of *Tbx1*, B12 acts on one or more target genes tht are also targeted by TBX1, perhaps by altering DNA or histone methylation, or other epigenetic mechanism. Alternatively, B12 might correct a metabolic abnormality caused by the loss of TBX1. Therefore, the possible mechanisms by which B12 acts in rodents are at least three, transcriptional, epigenetic and metabolic, as shown in the model in Fig. 4. Additional studies will be needed to elucidate which of the potential mechanisms are important for cortical development. Interestingly, a distinct metabolic profile has been reported for children with 22q11.2DS (Napoli et al., 2015), including abnormal levels of multiple metabolites in pathways that are B12-dependent.

How might TCP rescue the cortical phenotype in *Tbx1* mutants? Any proposed mechanism must be compatible with existing knowledge of TBX1 and LSD1 function in this context. For example, in the mouse, *Tbx1* is required in the embryonic medio-lateral (ML) cortex (precursor of the somatosensoty cortex) for the differentiation and migration of glutamatergic and GABAergic neurons, a role that is cell non-autonomous and mesoderm-dependent (Flore et al., 2017). From an epigenetic point of view, TBX1 has been shown to recruit histone methyltransferases (HMT) to primed enhancers, rather than to active enhancers, and at some chromatin sites it co-localizes with LSD1, although these sites have not been mapped. Mutation of *Tbx1* in cultured cells and *in vivo* was associated with reduced H3K4me1 at target loci, and this was rescued partially by TCP treatment, as was the cardiovascular phenotype in *Tbx1* mutant embryos (Fulcoli et al., 2016). We propose a model where TBX1, HMTs and LSD1 co-localize at one or more enhancers of an unknown mesodermally-expressed extracellular signal gene (ECS), activating its expression and thereby promoting appropriate differentiation of cortical progenitors in the ML-cortex. In *Tbx1* mutant embryos, the reduced presence of TBX1 and HMTs at the enhancer/s, while maintaining LSD1, reduces levels of H3K4me1 locally causing, or contributing to, a repressive chromatin environment, and silencing of the ECS gene. TCP treatment, by blocking LSD1, counteracts the reduced H3K4 methylation caused by *Tbx1* mutation and restores expression of the ECS gene and thereby appropriate neuronal differentiation.

TCP and B12 can now be added to a growing list of drugs and compounds with proven ability to ameliorate the neuronal and/or behavioral deficits found in mouse models of 22q11.2DS (Armando et al., 2020; Fernandez et al., 2019; Mukherjee et al., 2019; Nilsson et al., 2018; Tamura et al., 2016) and, in one case, in patients with 22q11.2DS (Armando et al., 2020). To our knowledge, this study is the first to identify drugs that target *TBX1*, the major candidate gene for 22q11.2DS, that have preventative potential in the mouse model. TCP is not an ideal therapeutic agent because it lacks specificity for LSD1; it was originally approved by the US Food and Drug Administration (FDA) for the treatment of depression based on its ability to inhibit monoamine oxidase (MAO). However, numerous TCP derivatives have been developed that have greater specificity for LSD1, some of which are in clinical trials for cancer therapy, reviewed in (Fang et al., 2019). B12 is an exciting therapeutic candidate because it is an approved drug, including for prenatal use, that is inexpensive and virtually harmless, even at high doses. If proven effective in the clinical setting, it would be an important advance over current therapeutic options, which are limited to treating symptoms. In the mean time, mouse studies will continue to provide critical information about the optimal timing of B12 treatment, or indeed of any other therapy, given that we do not know whether the brain-related abnormalities in 22q11.2DS patients (or in the mouse model) are preventable using post-natal treatment, or whether they are the irreversible consequences of abnormal prenatal development. Finally, given the proven preventive potential of high dose B12 treatment in the mouse, biochemical analysis of patients would be appropriate to determine whether they have B12 deficiency or in B12-dependent pathways.

## AUTHOR CONTRIBUTIONS

E.I. and A.B. designed the study. I.F. and G.F. performed the experiments. S.C. and G.L. assisted with experiments. E.I., I.F. and G.F. analyzed the data. E.I. wrote the manuscript.

## ACKNOWLEDGEMENTS

We are grateful for the invaluable support of the staff of the IGB-CNR mouse facility and the Integrated Microscopy facility. The study was supported by grants from the Jerome Lejeune Foundation, project No. 1685 (to E.I.), the Leducq Foundation Transatlantic Network of Excellence in Cardiovascular Research, 15CVD01 (to E.I. and A.B.) and the Ministero dell’Istruzione dell’Università e della Ricerca (MUIR), #20179J2P9J (to E.I. and A.B).

## CONFLICT OF INTEREST

None

## Notes

### Competing Interest Statement

The authors have declared no competing interest.

## REFERENCES

Armando, M., Ciampoli, M., Padula, M. C., Amminger, P., De Crescenzo, F., Maeder, J., Schneider, M., Schaer, M., Managò, F., Eliez, S., et al. (2020). Favorable effects of omega-3 polyunsaturated fatty acids in attentional control and conversion rate to psychosis in 22q11.2 deletion syndrome. Neuropharmacology 168, 107995.

Binda, C., Mattevi, A. and Edmondson, D. E. (2002). Structure-function relationships in flavoenzyme-dependent amine oxidations: a comparison of polyamine oxidase and monoamine oxidase. J. Biol. Chem. 277, 23973–23976.

Braunstein, E. M., Monks, D. C., Aggarwal, V. S., Arnold, J. S. and Morrow, B. E. (2009). Tbx1 and Brn4 regulate retinoic acid metabolic genes during cochlear morphogenesis. BMC Dev. Biol. 9, 31.

Calmont, A., Ivins, S., Van Bueren, K. L., Papangeli, I., Kyriakopoulou, V., Andrews, W. D., Martin, J. F., Moon, A. M., Illingworth, E. A., Basson, M. A., et al. (2009). Tbx1 controls cardiac neural crest cell migration during arch artery development by regulating Gbx2 expression in the pharyngeal ectoderm. Dev. Camb. Engl. 136, 3173–3183.

Caprio, C. and Baldini, A. (2014). p53 suppression partially rescues the mutant phenotype in mouse models of DiGeorge syndrome. Proc. Natl. Acad. Sci. 201401923.

Fagman, H., Liao, J., Westerlund, J., Andersson, L., Morrow, B. E. and Nilsson, M. (2007). The 22q11 deletion syndrome candidate gene Tbx1 determines thyroid size and positioning. Hum. Mol. Genet. 16, 276–285.

Fang, Y., Liao, G. and Yu, B. (2019). LSD1/KDM1A inhibitors in clinical trials: advances and prospects. J. Hematol. Oncol.J Hematol Oncol 12, 129.

Fernandez, A., Meechan, D. W., Karpinski, B. A., Paronett, E. M., Bryan, C. A., Rutz, H. L., Radin, E. A., Lubin, N., Bonner, E. R., Popratiloff, A., et al. (2019). Mitochondrial Dysfunction Leads to Cortical Under-Connectivity and Cognitive Impairment. Neuron 102, 1127–1142.e3.

Flore, G., Cioffi, S., Bilio, M. and Illingworth, E. (2017). Cortical Development Requires Mesodermal Expression of Tbx1, a Gene Haploinsufficient in 22q11.2 Deletion Syndrome. Cereb. Cortex N. Y. N 1991 27, 2210–2225.

Forneris, F., Binda, C., Vanoni, M. A., Mattevi, A. and Battaglioli, E. (2005). Histone demethylation catalysed by LSD1 is a flavin-dependent oxidative process. FEBS Lett. 579, 2203–2207.

Fulcoli, F. G., Franzese, M., Liu, X., Zhang, Z., Angelini, C. and Baldini, A. (2016). Rebalancing gene haploinsufficiency in vivo by targeting chromatin. Nat Commun 7, 11688.

Green, R., Allen, L. H., Bjørke-Monsen, A.-L., Brito, A., Guéant, J.-L., Miller, J. W., Molloy, A. M., Nexo, E., Stabler, S., Toh, B.-H., et al. (2017). Vitamin B12 deficiency. Nat. Rev. Dis. Primer 3, 17040.

Haddad, R. A., Clines, G. A. and Wyckoff, J. A. (2019). A case report of T-box 1 mutation causing phenotypic features of chromosome 22q11.2 deletion syndrome. Clin. Diabetes Endocrinol. 5, 13.

Hirsch, M. R., Tiveron, M. C., Guillemot, F., Brunet, J. F. and Goridis, C. (1998). Control of noradrenergic differentiation and Phox2a expression by MASH1 in the central and peripheral nervous system. Dev. Camb. Engl. 125, 599–608.

Lania, G., Zhang, Z., Huynh, T., Caprio, C., Moon, A. M., Vitelli, F. and Baldini, A. (2009). Early thyroid development requires a Tbx1-Fgf8 pathway. Dev. Biol. 328, 109–117.

Lania, G., Bresciani, A., Bisbocci, M., Francone, A., Colonna, V., Altamura, S. and Baldini, A. (2016). Vitamin B12 ameliorates the phenotype of a mouse model of DiGeorge syndrome. Hum. Mol. Genet.

Lindsay, E. A., Vitelli, F., Su, H., Morishima, M., Huynh, T., Pramparo, T., Jurecic, V., Ogunrinu, G., Sutherland, H. F., Scambler, P. J., et al. (2001). Tbx1 haploinsufficieny in the DiGeorge syndrome region causes aortic arch defects in mice. Nature 410, 97–101.

Motahari, Z., Moody, S. A., Maynard, T. M. and LaMantia, A.-S. (2019). In the line-up: deleted genes associated with DiGeorge/22q11.2 deletion syndrome: are they all suspects? J. Neurodev. Disord. 11, 7.

Mukherjee, A., Carvalho, F., Eliez, S. and Caroni, P. (2019). Long-Lasting Rescue of Network and Cognitive Dysfunction in a Genetic Schizophrenia Model. Cell 178, 1387–1402.e14.

Napoli, E., Tassone, F., Wong, S., Angkustsiri, K., Simon, T. J., Song, G. and Giulivi, C. (2015). Mitochondrial Citrate Transporter-dependent Metabolic Signature in the 22q11.2 Deletion Syndrome. J. Biol. Chem. 290, 23240–23253.

Nilsson, S. R. O., Heath, C. J., Takillah, S., Didienne, S., Fejgin, K., Nielsen, V., Nielsen, J., Saksida, L. M., Mariani, J., Faure, P., et al. (2018). Continuous performance test impairment in a 22q11.2 microdeletion mouse model: improvement by amphetamine. Transl. Psychiatry 8, 247.

Ogata, T., Niihori, T., Tanaka, N., Kawai, M., Nagashima, T., Funayama, R., Nakayama, K., Nakashima, S., Kato, F., Fukami, M., et al. (2014). TBX1 mutation identified by exome sequencing in a Japanese family with 22q11.2 deletion syndrome-like craniofacial features and hypocalcemia. PloS One 9, e91598.

Paylor, R. and Lindsay, E. (2006). Mouse models of 22q11 deletion syndrome. Biol. Psychiatry 59, 1172–1179.

Paylor, R., Glaser, B., Mupo, A., Ataliotis, P., Spencer, C., Sobotka, A., Sparks, C., Choi, C.-H., Oghalai, J., Curran, S., et al. (2006). Tbx1 haploinsufficiency is linked to behavioral disorders in mice and humans: implications for 22q11 deletion syndrome. Proc. Natl. Acad. Sci. U. S. A. 103, 7729–7734.

Prasad, S. E., Howley, S. and Murphy, K. C. (2008). Candidate genes and the behavioral phenotype in 22q11.2 deletion syndrome. Dev. Disabil. Res. Rev. 14, 26–34.

Racedo, S. E., Hasten, E., Lin, M., Devakanmalai, G. S., Guo, T., Ozbudak, E. M., Cai, C.-L., Zheng, D. and Morrow, B. E. (2017). Reduced dosage of β-catenin provides significant rescue of cardiac outflow tract anomalies in a Tbx1 conditional null mouse model of 22q11.2 deletion syndrome. PLoS Genet. 13, e1006687.

Shi, Y., Lan, F., Matson, C., Mulligan, P., Whetstine, J. R., Cole, P. A., Casero, R. A. and Shi, Y. (2004). Histone demethylation mediated by the nuclear amine oxidase homolog LSD1. Cell 119, 941–953.

Tamura, M., Mukai, J., Gordon, J. A. and Gogos, J. A. (2016). Developmental Inhibition of Gsk3 Rescues Behavioral and Neurophysiological Deficits in a Mouse Model of Schizophrenia Predisposition. Neuron 89, 1100–1109.

Torres-Juan, L., Rosell, J., Morla, M., Vidal-Pou, C., García-Algas, F., de la Fuente, M.-A., Juan, M., Tubau, A., Bachiller, D., Bernues, M., et al. (2007). Mutations in TBX1 genocopy the 22q11.2 deletion and duplication syndromes: a new susceptibility factor for mental retardation. Eur. J. Hum. Genet. EJHG 15, 658–663.

Vitelli, F., Lania, G., Huynh, T. and Baldini, A. (2010). Partial rescue of the Tbx1 mutant heart phenotype by Fgf8: genetic evidence of impaired tissue response to Fgf8. J Mol Cell Cardiol 49, 836–40.

Xu, H., Morishima, M., Wylie, J. N., Schwartz, R. J., Bruneau, B. G., Lindsay, E. A. and Baldini, A. (2004). Tbx1 has a dual role in the morphogenesis of the cardiac outflow tract. Dev. Camb. Engl. 131, 3217–3227.

Xu, H., Chen, L. and Baldini, A. (2007). In vivo genetic ablation of the periotic mesoderm affects cell proliferation survival and differentiation in the cochlea. Dev. Biol. 310, 329–340.

Zhang, Z., Cerrato, F., Xu, H., Vitelli, F., Morishima, M., Vincentz, J., Furuta, Y., Ma, L., Martin, J. F., Baldini, A., et al. (2005). Tbx1 expression in pharyngeal epithelia is necessary for pharyngeal arch artery development. Dev. Camb. Engl. 132, 5307–5315.

